# vafSOMA: Detecting cell-layer-fixed somatic mutations in plants

**DOI:** 10.64898/2026.11.02.763622

**Authors:** Ming Zhou, Frank Johannes

**Affiliations:** Plant Epigenomics, Technical University of Munich, Emil-Ramann-Str. 4, 85354 Freising, Germany; Centre for Research in Agricultural Genomics (CRAG), CSIC-IRTA-UAB-UB, Campus UAB, Bellaterra, Barcelona, Spain

**Keywords:** somatic mutation, shoot apical meristem, histogenic layers, periclinal chimera, somatic mutation rate, variant allele frequency, mixture model, Mutect2, long-lived plants

## Abstract

Somatic mutations are a source of heritable genetic variation in plants and have long been exploited by breeders through selection and vegetative propagation of bud sports. They often arise in specific meristem layers and can become fixed as descendant cell lineages expand during development. Bulk sequencing of plant organs mixes DNA from these layers, so the expected variant allele frequency of a layer-fixed mutation depends on that layer’s tissue contribution, which complicates their detection. Here we present `vafSOMA`, a statistical model that uses tissue composition to detect fixed somatic mutations, assign them to layers and obtain counts for mutation-rate estimation. It accommodates bulk and layer-enriched samples, including those that retain DNA from other layers after enrichment. In simulations, `vafSOMA` performed close to the best variant allele frequency cutoff chosen in hindsight, without requiring a user-selected cutoff. Its diagnostics flagged layers for which the data did not support separate mutation counts or rates. Analysing replicate organs substantially improved detection of mutations fixed in the branch meristem and yielded estimates close to the simulated stem-cell mutation rate. Application to apricot sequencing data showed that most reference mutations fixed in the dominant layer were recovered from fruit mixtures at high depth, whereas epidermal mutations remained unresolvable in fruit and leaves. These findings suggest that published bulk somatic sequencing studies mainly capture mutations fixed in the dominant tissue layer and may therefore underestimate the combined input of fixed mutations by roughly two- to threefold when layer-specific rates are similar. `vafSOMA` is available at https://github.com/jlab-code/vafSOMA.

## Introduction

Somatic mutations generate genetic differences among branches and organs as plants grow. New leaves, flowers and branches arise from shoot apical meristems (SAMs), populations of stem cells located at the tip of each growing shoot (Lyndon, 1998). Because plants lack an early-segregated germline, mutations that accumulate in these stem cells can reach flowers and gametes, contributing to heritable variation in both clonally and sexually reproducing plants (Lanfear, 2018; Schoen and Schultz, 2019). In perennial fruit crops, somatic mutations occasionally produce bud sports, branches with altered fruit, flower or growth traits from which many cultivars have been derived through somatic selection and vegetative propagation (Foster and Aranzana, 2018; Ban and Jung, 2023). Such genetic mosaicism has also motivated ideas about selection among cell lineages and plant modules in natural populations (Whitham and Slobodchikoff, 1981; Gill et al., 1995; Otto and Hastings, 1998; Yu et al., 2020; Reusch et al., 2021).

The developmental fate of a mutation depends on its cell lineage. In most angiosperms, the SAM comprises three layers, L1 to L3, each maintained by a small population of apical stem cells. L1 produces the epidermis, L2 the subepidermal tissues and, in many species, gametes, and L3 the inner tissues (Satina et al., 1940; Tilney-Bassett, 1986; Lyndon, 1998; Frank and Chitwood, 2016). These layers follow largely separate lineages, so a mutation can spread throughout a branch while remaining fixed in only one layer (Fig. 1c). Many bud sports are correspondingly layer-specific chimeras (Boss and Thomas, 2002; Franks et al., 2002; Vezzulli et al., 2012; Pelsy et al., 2015; Foster and Aranzana, 2018).

**Figure 1:**
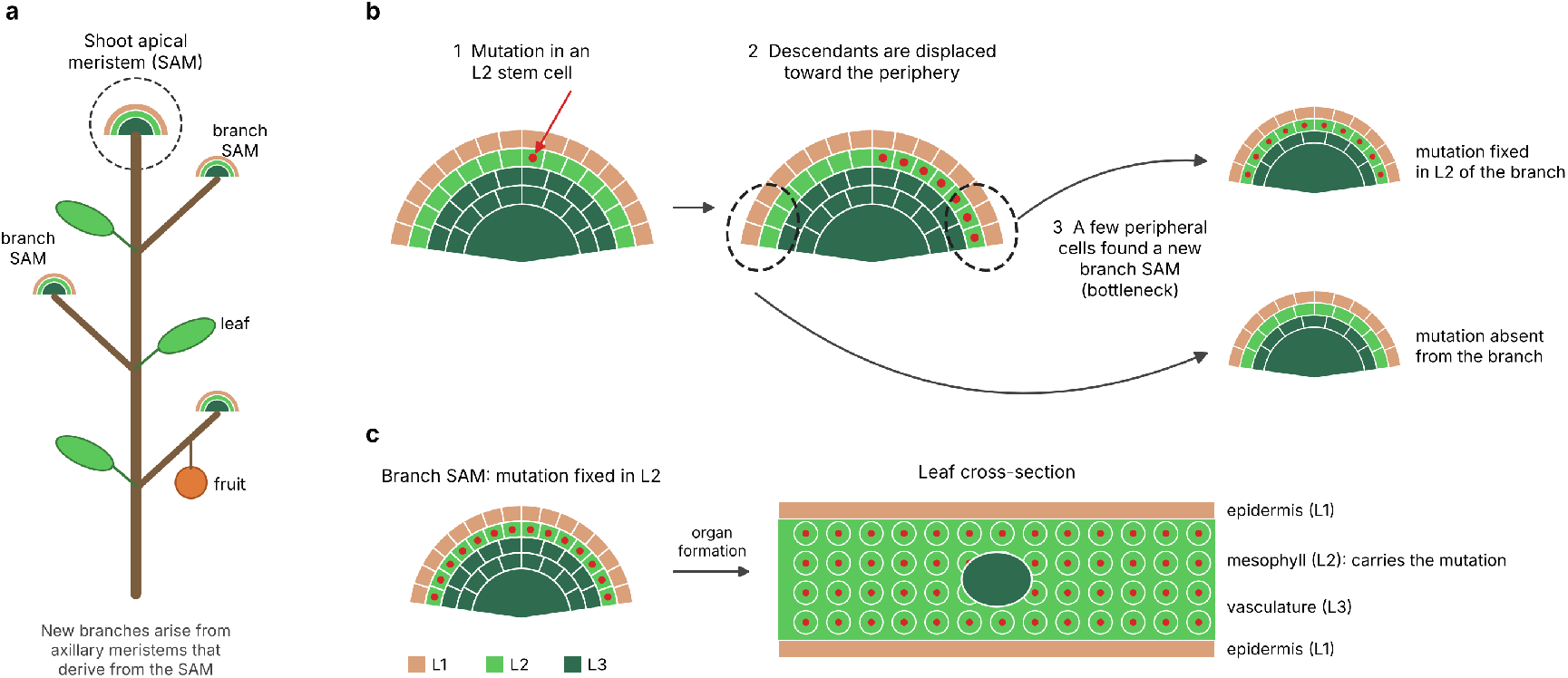
Origin and fixation of layer-specific somatic mutations. **a** Plants grow from shoot apical meristems (SAMs) at the tips of their shoots. Each SAM consists of three cell layers (L1, L2, L3), and each new branch arises from an axillary meristem derived from the parental SAM. **b** Side view of a SAM. (1) A mutation arises in one of the few stem cells at the summit of L2. (2) Its descendants are displaced toward the periphery, where they form a mutant clone. (3) A new branch SAM is founded from a few peripheral cells: if these derive from the mutant clone, the mutation becomes fixed in the L2 of the branch; otherwise it is absent. Repeated bottlenecks of this kind at branching and organ formation drive somatic mutations to fixation or loss. **c** Each layer is a separate cell lineage. A mutation fixed in the L2 of a branch SAM is carried by all L2-derived cells of the organs the branch produces, here the leaf mesophyll, but not by the L1-derived epidermis or the L3-derived vasculature. Organs are therefore periclinal chimeras; the consequences for bulk sequencing are shown in Fig. 2a.

A small group of slowly dividing stem cells at the centre of the SAM, the apical stem cells, maintains the SAM while producing descendants that proliferate toward its periphery. Small pools of these peripheral cells initiate new organs and axillary meristems, from which new branches develop (Lyndon, 1998; Burian et al., 2016; Shi et al., 2016) (Fig. 1a,b). If all founders of a layer carry a mutation, it can become fixed in that layer of the new branch meristem and pass into its descendant organs. Repeated branching therefore allows somatic drift to fix stem-cell mutations in large plant sectors (Burian et al., 2016; Tomimoto and Satake, 2023; Chen et al., 2024; Johannes, 2025a,b). Mutations arising later during organ development instead generate variation that remains largely confined to that organ (Johannes, 2026a).

Recent sequencing studies across diverse plant species have sought to detect somatic variants and estimate mutation rates (Xie et al., 2016; Schmid-Siegert et al., 2017; Plomion et al., 2018; Hanlon et al., 2019; Wang et al., 2019; Hofmeister et al., 2020; Orr et al., 2020; Shahryary et al., 2020; Duan et al., 2022; Pérez-Román et al., 2022; Satake et al., 2023; Zhou et al., 2025). These studies relied mainly on bulk sequencing of leaves. Relating these estimates to mutation accumulation along apical stem-cell lineages requires knowing which lineages the sampled organ represents and which mutations are counted as fixed (Johannes, 2025a).

A major challenge is that bulk sequencing mixes DNA from different layers, diluting the signal of mutations fixed in any one layer (Fig. 2a). In a diploid, a heterozygous mutation is further diluted by reads from the unmutated homolog. If layer *k* contributes a fraction *a*_*k*_ of the sequenced DNA, the expected variant allele frequency (VAF) of a mutation fixed in that layer is *a*_*k*_*/*2 against a haploid reference without phasing, or *a*_*k*_ when only reads phased to the mutated haplotype are counted (Goel et al., 2024; Amundson et al., 2025; Johannes, 2026a). In leaves, L2-derived mesophyll often dominates the sample, placing L2-fixed mutations at moderate VAFs. Mutations fixed in L1 and L3 occur at lower VAFs, where they overlap with non-fixed somatic variants and sequencing and mapping artefacts (Schmitt et al., 2022, 2024; Goel et al., 2024).

**Figure 2:**
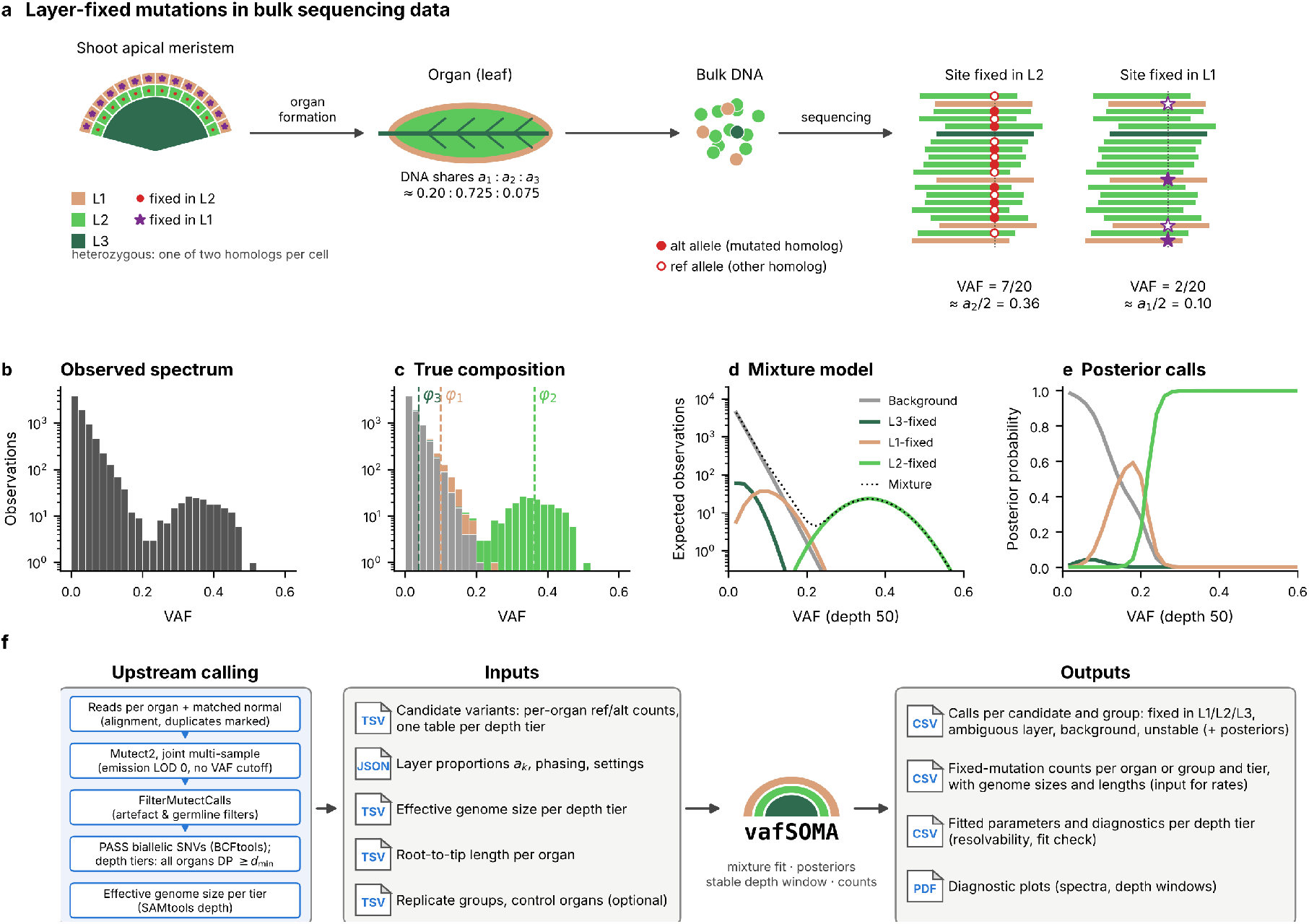
Overview of vafSOMA. **a** Mutations arising in shoot apical meristem stem cells become fixed within one layer (L1, L2 or L3). Organs derive from all three layers in unequal proportions. Because the mutation is heterozygous, every cell of the mutant layer carries it on one of its two homologs: in bulk DNA, reads from the mutated homolog carry the alternative allele (filled symbols) and reads from the other homolog the reference allele (open symbols), giving an expected VAF of *a*_*k*_*/*2 (unphased diploid). **b** VAF spectrum of candidate variants in simulated bulk data (eight organs, 50×, layer proportions 0.20*/*0.725*/*0.075). **c** The same spectrum coloured by true class; dashed lines mark the expected VAFs *φ*_*k*_. **d** Mixture components at depth 50× : the exponential background and the layer components, scaled by their weights. **e** Posterior probability of each class as a function of the observed VAF; these replace a fixed VAF cutoff. Minor-layer candidates reach only intermediate probabilities because their component overlaps the background. **f** Workflow: upstream candidate calling with Mutect2 (Materials and Methods), the inputs to `vafSOMA` and its outputs; the central symbol is the `vafSOMA` logo. In d and e, components are evaluated at the generating parameters for illustration.

Recent studies have shown that this dilution can be reduced by sequencing tissues enriched for individual layers, such as separated fruit peel and flesh (Goel et al., 2024) or isolated epidermal trichomes (Amundson et al., 2025). Analyses of these samples revealed that most detected somatic mutations were restricted to individual layers and that the epidermal lineage accumulated mutations faster than the inner layers (Goel et al., 2024; Amundson et al., 2025). However, enrichment depends on tissue accessibility and preparation and can remain incomplete. For example, in apricot, peel is only about 69% L1-derived (Goel et al., 2024).

Somatic variant detection in plants has improved with the adoption of cancer-derived callers such as Mutect2 (Benjamin et al., 2019) and Strelka2 (Kim et al., 2018), which compare samples with a matched normal to help distinguish somatic variants from germline variants and artefacts (Cibulskis et al., 2013; Schmitt et al., 2022). However, these tools do not model plant cell layers or distinguish mutations fixed within a layer from those confined to a subset of its cells. Plant studies commonly infer fixation by retaining candidates above an ad hoc VAF or alternative-read-count cutoff (Schmitt et al., 2022; Goel et al., 2024; Amundson et al., 2025). Raising this threshold removes more background but also loses fixed mutations from minor layers; lowering it retains more fixed mutations while admitting non-fixed variants and artefacts. Neither choice identifies the layer of fixation. An approach that accounts for layer contributions is therefore needed for both bulk and incompletely enriched samples.

To address this need, we present `vafSOMA`, which uses tissue composition to distinguish fixed somatic mutations from background and assign them to layers. It provides counts per organ or replicate group from which layer-specific mutation rates and confidence intervals can be estimated. We use cell-lineage simulations of plant development to test layer recovery, compare VAF cutoffs and assess the benefit of replicate organs for detection and rate estimation. We then analyse controlled mixtures of apricot fruit tissues and conventional leaf sequencing data to examine how these predictions translate to real sequencing data (Goel et al., 2024).

## Results

### Overview of vafSOMA

#### Input data

`vafSOMA` takes tables of candidate somatic variants, with reference and alternative read counts for every sampled organ (Fig. 2f). We generated these tables by jointly analysing aligned, duplicate-marked BAM files with Mutect2, using an early-diverging part of the same plant as the matched normal and applying no VAF cutoff (Materials and Methods). After filtering, we retained biallelic single-nucleotide variants. Each table records read counts, depth and VAF in every organ, including observations with no alternative reads. Separate tables are supplied for each minimum-depth cutoff, together with the corresponding effective genome size, defined as the number of callable haploid sites.

The user supplies the sequencing design (bulk or layer-enriched), tissue composition, phasing, ploidy and sample ages or path lengths. Tissue contributions can be estimated from published data or independent measurements, and organs can be assigned to replicate groups where available. Although tested only with Mutect2 output, the model can accept read-count tables from other callers in the required format.

#### Fixed variant detection and cell-layer assignment

`vafSOMA` models candidate read counts as a mixture of mutations fixed in individual layers and a background of non-fixed somatic variants and residual artefacts (Fig. 2c,d). For a single organ, the basic binomial model is

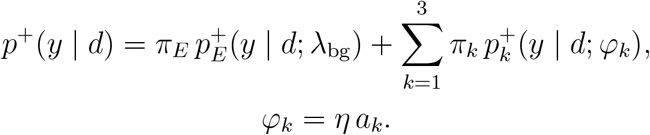

Here, *y* is the alternative read count, *d* the total depth, and the mixture weights satisfy 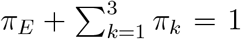. Each layer component describes read counts with an expected VAF *φ*_*k*_, determined by that layer’s DNA contribution *a*_*k*_. The factor *η* is 1*/*2 for unphased and 1 for phased counts in a diploid. The superscript + denotes conditioning on at least one alternative read. The read model can also incorporate sequencing error and beta-binomial overdispersion (Materials and Methods).

Tissue composition therefore determines the expected VAFs of fixed mutations, while the mixture weights and background shape are estimated from the data. Posterior probabilities quantify support for each candidate being background or fixed in a particular layer (Fig. 2e).

#### Choice of background model

We used low-VAF candidates from the layer-enriched apricot fruit samples of Goel et al. (2024) to guide the background model. Enrichment shifts mutations fixed in the dominant layer to higher VAFs, allowing the background below them to be examined separately. We found that this background was unimodal and slightly lighter-tailed than exponential. An exponential distribution nevertheless provided a conservative approximation (Materials and Methods; File S3, Figure S1).

We therefore modelled the underlying background VAF *q* using the truncated exponential density

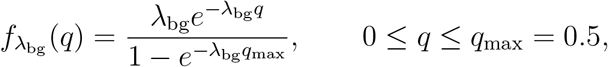

where *λ*_bg_ controls the decay of the background. The corresponding read-count component 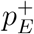 accounts for sampling at the observed depth and conditioning on detection.

#### Replicate modelling

Organs from the same branch can be analysed jointly as a replicate group. They should be sampled close together along the branch and have formed close in time. A mutation fixed in their shared branch meristem is expected in every replicate, whereas a non-fixed variant may be confined to one organ. The model combines read counts across replicates under a common fixation state, while allowing both shared and organ-specific background variation. This helps distinguish fixation inferred from one organ from fixation supported across several. With one organ per group, the model reduces to the single-organ model (Materials and Methods).

#### All-organs screen

`vafSOMA` can optionally remove candidates with alternative reads in every non-control organ before fitting. This screen is intended for sampling designs with no branch shared by all non-control organs after divergence from the matched normal. Candidates with this pattern often reflect recurrent artefacts, such as mapping errors, and can distort the estimated background. Screening them before fitting limits this influence. The screen can also exclude rare genuine mutations; its assumptions and limitations are described in Materials and Methods.

#### Diagnostics and layer resolvability

`vafSOMA` assesses both support for individual layers and agreement with the overall VAF spectrum. The model-implied recall measures how often a mutation fixed in a layer would be assigned correctly under the fitted model. A layer is flagged as unresolvable if this recall is below 0.5 or its expected VAF lies within the fitted background. A separate fit check compares the observed and fitted spectra and raises a warning for substantial discrepancies. Neither diagnostic changes fitted posteriors or individual calls. An unresolvable flag indicates insufficient support for a separate layer-specific count or rate.

#### Mutation-rate estimation

Posterior-weighted counts include every candidate according to its probability of fixation in each layer, including candidates without a confident layer assignment. For a resolvable layer, these counts yield a rate per site and haploid genome per unit of length or time,

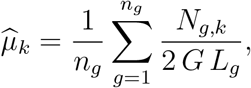

where *N*_*g,k*_ is the posterior-weighted fixed-mutation count for layer *k* in organ or replicate group *g, G* is the effective haploid genome size, and 2*G* is the diploid mutational target. *L*_*g*_ is the age or path length since divergence from the matched normal, and *n*_*g*_ is the number of non-control groups. `vafSOMA` reports counts for each minimum-depth cutoff and selects a stable window for summarising the results. We calculated rates from these outputs in a separate step, averaging across the selected window and obtaining bootstrap confidence intervals (Materials and Methods).

### Simulation study

#### Simulation setup

To test mutation recovery against known fixation states, we generated synthetic data with simSOMA, which tracks cell lineages and somatic mutations through plant development (Johannes, 2026a,b). We simulated the approximate branching topology of the apricot tree studied by Goel et al. (2024), with a leaf and a fruit at each of seven branch tips.

Each layer evolved independently, producing mutations across the full range of frequencies within organs. Non-fixed variants therefore arose from the simulated developmental process. We calculated bulk VAFs from the fraction of cells carrying each mutation within its layer and that layer’s DNA contribution to the sampled organ.

We then generated read counts using an observation model calibrated on the apricot input tables, incorporating variation in depth, sequencing error, overdispersion and selection during candidate calling (File S1, Section S7). Background candidates were resampled from the corresponding apricot data after excluding sites with evidence of real somatic mutations in the layer-enriched samples. The stem-cell, developmental and observation-model parameters are given in Materials and Methods.

#### Variant recovery from bulk data depends on layer composition

We varied layer contributions while keeping the developmental model unchanged. The L1 share ranged from 0.05 to 0.45, L3 was fixed at 0.075, and L2 contributed the remainder (Fig. 3; File S2, Tables S1 and S2). At this contribution, L3’s expected VAF was about 0.04 and overlapped the background. Including an L3 component attributed background variation to L3 fixation, so we fitted only L1 and L2 (Materials and Methods). For these two layers, we calculated counts by summing candidate fixation probabilities, including uncertain assignments, and reported them only where the layer was classified as resolvable.

**Figure 3:**
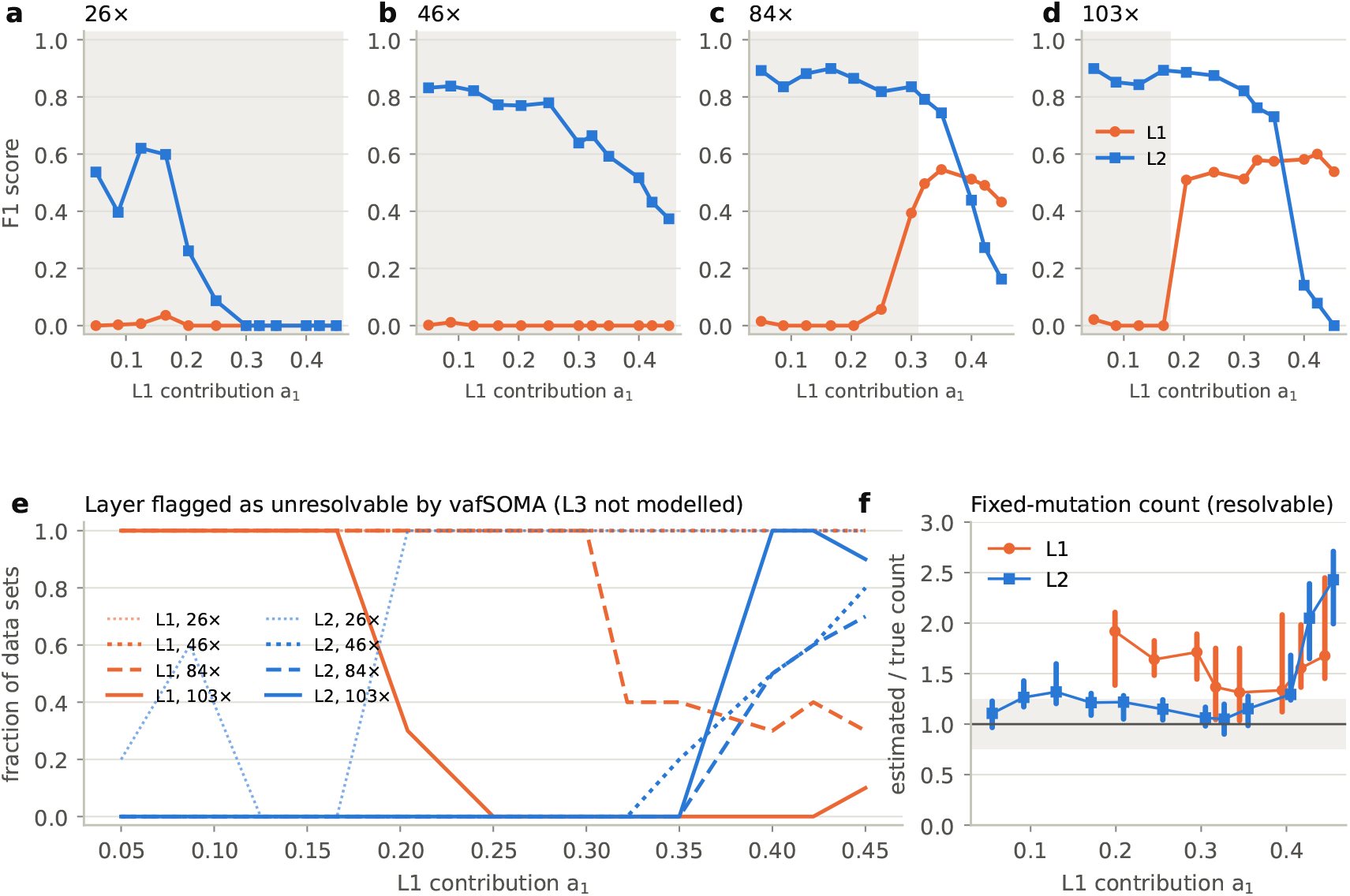
What bulk data can resolve depends on layer composition. simSOMA trees (apricot topology, fruit designs with six non-normal fruits), L1 contribution *a*_1_ from 0.05 to 0.45, *a*_3_ = 0.075, *a*_2_ = 0.925 − *a*_1_; reads from the observation model calibrated on the apricot designs; ten trees per setting; `vafSOMA’s` per-organ model with the all-organs screen, L1 and L2 modelled. **a–d** Mean F1 score for mutations fixed in L1 and L2 as a function of *a*_1_ at 26, 46, 84 and 103× (depth at true mutations); shading, compositions at which L1 is flagged as unresolvable in at least half of the data sets. **e** Fraction of data sets in which `vafSOMA` flags L1 or L2 as unresolvable, by depth. **f** Posterior-weighted fixed-mutation count relative to the truth for L1 and L2 in data sets where the layer is resolvable (median and interquartile range over trees and depths; grey band, ±25%); depth-aware fixation.

L2 was recovered most reliably when it clearly dominated the sample. At depths of 46–103×, its mean F1 score was 0.77–0.90 for L1 shares up to 0.25, falling to 0.64–0.84 at 0.30. Median fixed-mutation counts were 1.03–1.34 times the truth. At 26×, L2 was flagged as unresolvable from an L1 share of 0.20.

L1 recovery depended on its separation from both the background and L2. At contributions of 0.17 or less, its expected VAF lay within the background, and `vafSOMA` made essentially no L1 calls and flagged the layer as unresolvable at every tested depth. At 103×, L1 was reported as resolvable in 70% of data sets at a contribution of 0.20 and in 90–100% from 0.25. At 84×, it was resolvable in 60–70% of data sets from a contribution of 0.32; at lower depths, it remained unresolvable. Where L1 was resolvable, its mean F1 score was 0.43–0.60. Counts were close to the truth at 84× (medians 1.01–1.05 at L1 contributions of 0.32– 0.40), but reached 1.6–1.9 times the truth at 103× (Fig. 3f).

As the contribution of L1 increased further, L2’s contribution fell and their expected VAFs converged. At L1 shares of 0.40 or more, this overlap made layer assignment increasingly uncertain. A candidate could therefore be detected as fixed without a confident assignment to L1 or L2.

#### vafSOMA performs close to the best VAF cutoff without tuning

A common alternative to modelling the VAF spectrum is to retain candidates above a fixed cutoff. We compared `vafSOMA` with cutoffs of 0.15, 0.20 and 0.25 across four compositions and depths of 26–103× (160 data sets; Fig. 4; File S2, Table S3). Because the true mutations were known, we could identify the cutoff that maximised the F1 score in each data set. This optimum ranged from 0.08 to 0.40 with composition and depth (Fig. 4a). Consequently, a cutoff that worked well in one setting could perform poorly in another. For a cutoff of 0.25, the mean F1 loss relative to the optimum was 0.17 and 0.21 at L1 shares of 0.32 and 0.42; for a cutoff of 0.15, it was 0.24 at an L1 share of 0.09.

**Figure 4:**
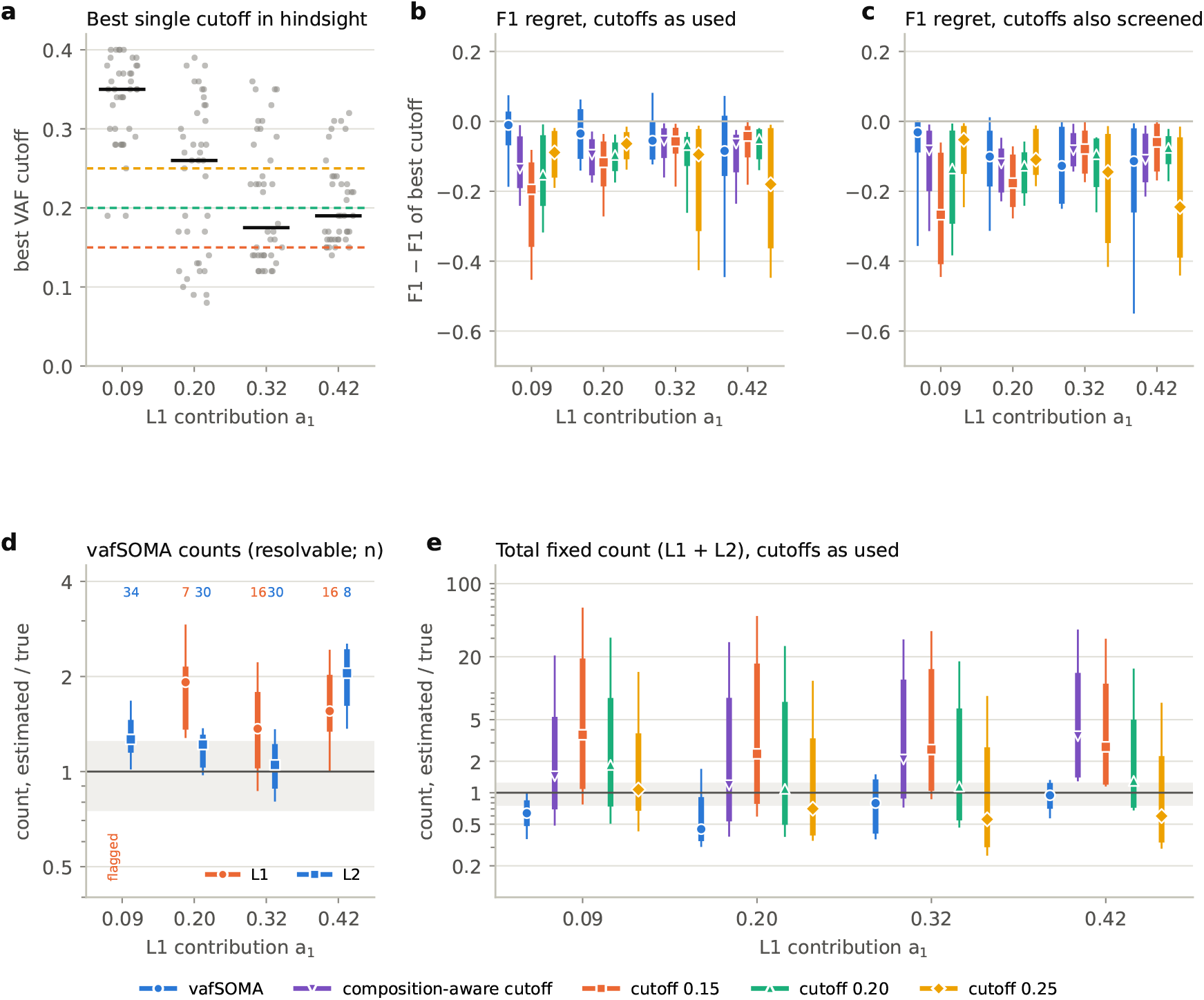
Comparison with fixed VAF cutoffs on simulated apricot trees. simSOMA trees, observation model calibrated on the apricot designs; L1 contribution *a*_1_ (L3 0.075, L2 the remainder); fruit designs at 26–103×; ten trees per setting; `vafSOMA` 0.4.2 with the all-organs screen. **a** Cutoff that maximizes the F1 score in each data set (tree × depth), by composition; bars, medians; dashed lines, the three fixed cutoffs. **b** F1 score relative to that best cutoff for `vafSOMA` (with unresolvable layers removed), a composition-aware cutoff (midway between a background VAF of 0.02 and the expected L2 VAF *φ*_2_) and the three fixed cutoffs, as commonly applied; points, medians; thick bars, interquartile ranges; thin bars, 10–90% ranges. **c** As **b**, with the all-organs screen applied to the cutoffs as well and the regret taken relative to the best screened cutoff. **d** Posterior-weighted counts of `vafSOMA` for L1 and L2 relative to the truth, for layers not flagged as unresolvable (log scale; grey band, ±25%; numbers, data sets). **e** Total count of fixed L1 and L2 mutations relative to the truth: layer-agnostic counts of the cutoffs and the sum of the resolvable layers of `vafSOMA` (where L1 is flagged, the `vafSOMA` total covers L2 only). Depth-aware fixation; strict fixation: File S3, Figure S2.

`vafSOMA` required no cutoff selection and had mean F1 losses of 0.02–0.11 across compositions (0.054 overall), compared with overall losses of 0.11–0.14 for the three fixed cutoffs (Fig. 4b). We also tested a cutoff calculated from tissue composition, placed halfway between a typical background VAF of 0.02 and the expected L2 VAF. Its mean loss was 0.100. Repeating the comparison with a stricter definition of fixation gave similar results (File S3, Figure S2).

Part of this difference reflected the all-organs screen, which was applied to `vafSOMA` but not to the cutoffs above. Applying the same screen to the cutoffs improved their F1 scores. With the best screened cutoff as the reference, mean F1 losses were similar: 0.13 for `vafSOMA`, 0.13–0.15 for the fixed cutoffs and 0.12 for the composition-based cutoff (Fig. 4c). With comparable screening, the methods therefore achieved similar calling accuracy on average, although the accuracy of the 0.15 and 0.25 cutoffs depended more strongly on composition (mean losses 0.08–0.25 across compositions, compared with 0.09–0.19 for `vafSOMA`; File S2, Table S3). `vafSOMA` additionally provided layer assignments, posterior-weighted counts, diagnostics of layer support and joint analysis of replicate organs.

We also examined fixed-mutation counts because these determine mutation-rate estimates (Fig. 4d,e). For resolvable L2, the median `vafSOMA` count was 1.22 times the truth, with 55% of estimates within 25% of the true count. For resolvable L1, the median was 1.54, with 23% within 25% (Fig. 4d). A VAF cutoff yields a combined count without layer assignment. Relative to the true total of fixed L1 and L2 mutations, cutoffs of 0.15, 0.20 and 0.25 gave median ratios of 3.05, 1.30 and 0.75, respectively. These ratios depended strongly on composition, showing that a fixed cutoff did not provide a consistent estimate of mutation counts across mixtures (Fig. 4e).

When L1 was unresolvable, `vafSOMA` reported only an L2 count, so this number could not be interpreted as the total across layers. This distinction mattered in these simulations: L1 accumulated mutations faster than L2, and L2 accounted for only about a third of the fixed mutations. Even a correct L2 estimate therefore described only part of the simulated mutation accumulation. `vafSOMA` made this limitation explicit by flagging L1 as unresolvable. A fixed VAF cutoff gave a single count without indicating which layers it represented or whether mutations fixed in other layers had been missed.

#### Diagnostics assess layer support and model fit

`vafSOMA` supplements its layer-specific counts with two diagnostics (Materials and Methods). The first assesses whether mutations fixed in each modelled layer can be distinguished from the background and other layers, flagging unsupported layers as unresolvable. The second compares the observed and fitted VAF spectra and raises a fit warning for substantial discrepancies.

We tested changes in the number and VAF distribution of background candidates, increased read-count overdispersion or sequencing error, and misspecified the L1 share by ±25% (Fig. 5; File S2, Tables S4 and S5). L2 counts changed little. Their median was 1.22 times the truth under the reference background and 1.13– 1.24 under the other conditions, except when the assumed L1 share was too high (1.37). The combined L1 and L2 count from a 0.20 cutoff was more sensitive to the background, rising from 1.30 times the truth in the reference setting to 1.93 with twice as many background candidates and 7.1 with the heavier-tailed component (Fig. 5a). Minor-layer counts required greater caution: for resolvable L1, median counts were 1.4–1.8 times the truth under most stress conditions (Fig. 5b).

**Figure 5:**
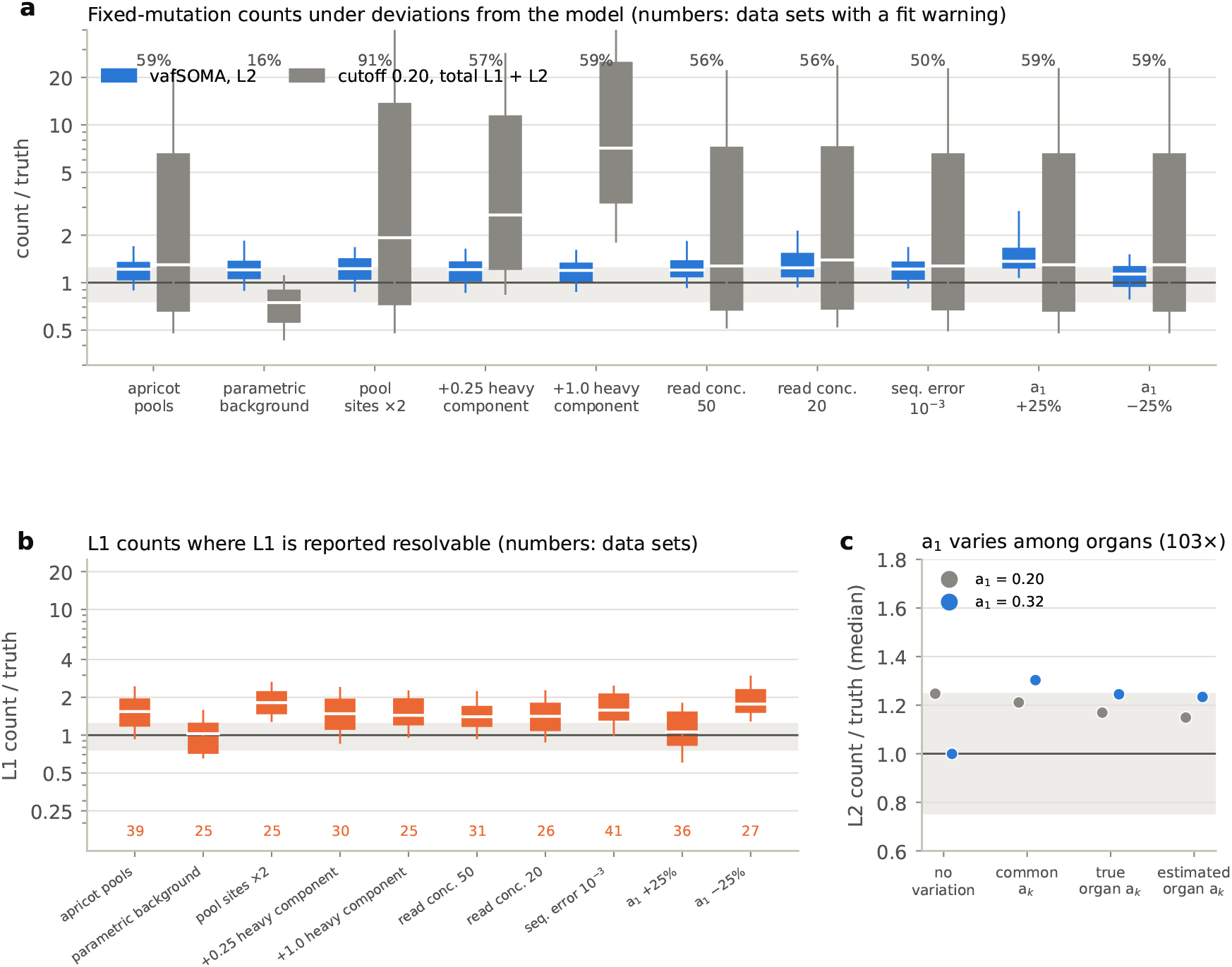
Diagnostics and robustness. simSOMA trees (apricot topology), observation model calibrated on the apricot designs, `vafSOMA` 0.4.2 per-organ model with the all-organs screen, depth-aware fixation rule. **a** Posterior-weighted L2 count of `vafSOMA` (resolvable L2 only) and total count of a 0.20 cutoff (L1 and L2), each relative to the truth, under the resampled apricot background (reference), a parametric background, twice the background pool, an additional heavy background component (0.25 or 1.0 times the pool size), read concentrations of 50 and 20 (reference 140), a sequencing error of 10^*−*3^ (reference 3 ×10^*−*4^) and an L1 contribution misspecified by 25%, over all compositions (*a*_1_ = 0.09–0.42) and depths (26–103×; 160 data sets per setting). Boxes, quartiles; lines, 10th–90th percentiles; shaded, ±25%. **b** Posterior-weighted L1 count of `vafSOMA` relative to the truth in the data sets in which L1 is reported resolvable (numbers). **c** Organ-to-organ variation in the L1 contribution (about ±50%) at 103×: median L2 count without variation and, with variation, for the common nominal composition, the true organ-specific compositions and compositions estimated from first-pass L2 calls (ten trees). Numbers above the boxes in **a**, fraction of data sets with a fit warning.

The diagnostic tests clarified how these outputs should be interpreted. Assessing both model-implied recall and separation from the background identified cases in which background candidates otherwise appeared to support a minor-layer component. We used these criteria to determine which layers supported separate count estimates. Fit warnings identified discrepancies in the overall spectrum but did not consistently track count bias. Variation in composition among organs could also affect counts without triggering a warning (Fig. 5c). Detailed evaluations of background overlap, fit warnings, depth and organ-specific composition are presented in Appendix B and File S2, Tables S4–S6. Together, these diagnostics provide quantitative checks on layer support and model fit that a fixed VAF cutoff alone does not offer.

#### Replicate organs improve detection of fixed variants

Mutations confined to part of an organ can have VAFs close to those expected for fixation within a layer. Evidence from several organs formed close in time can help distinguish these variants from mutations fixed in their shared branch meristem.

We simulated three leaves formed about one week apart at each branch tip and analysed one, two or three leaves. Single-leaf analysis overestimated the number of mutations fixed in the branch meristem by about 40% (median 1.38 times the truth). Joint analysis of two or three leaves reduced this excess, giving L2 counts of 1.18 and 1.00 times the truth, respectively (Fig. 6c; File S2, Table S7). F1 scores against meristem fixation increased from 0.75–0.78 with one leaf to 0.86–0.90 with two and 0.82–0.95 with three (Fig. 6a,b). With three leaves, the replicate-group model gave the best L2 calls in every setting. With two, it performed similarly to a simple concordance screen requiring a call in both leaves, but retained more meristem-fixed mutations (sensitivity 0.97 versus 0.84).

**Figure 6:**
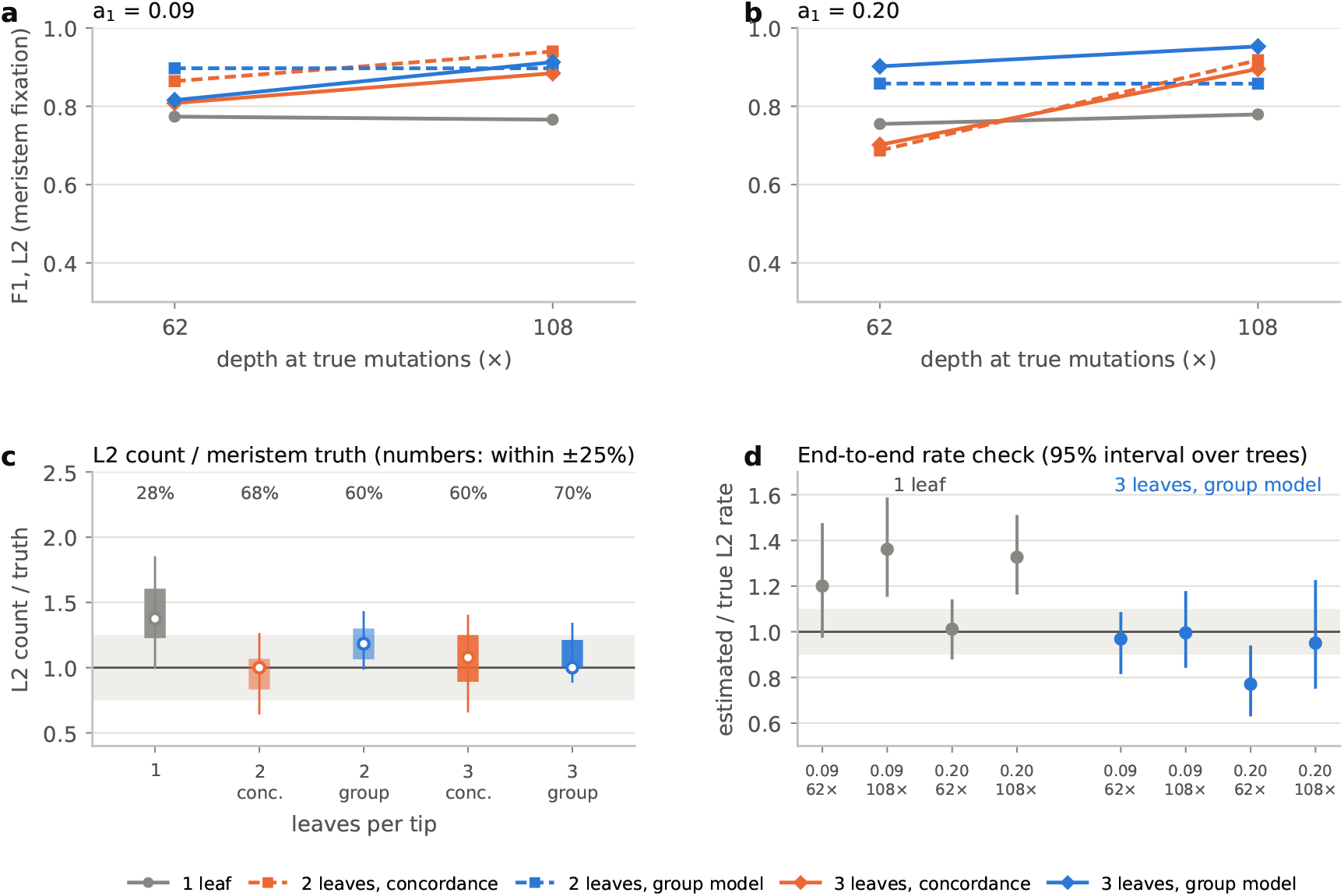
Replicate organs improve the calling of fixed mutations. simSOMA trees with three leaves per branch tip formed about one week apart (apricot topology, observation model calibrated on the apricot leaf designs), analysed with one leaf per tip (per-organ model), or with two or three leaves per tip (per-organ model with a concordance screen requiring a fixed call in at least two leaves; replicate-group model); `vafSOMA` 0.4.2 with the all-organs screen. Truth, fixation in the branch meristem (all leaves of the tip). **a**,**b** F1 score of L2 calls by depth at the true mutations at *a*_1_ = 0.09 and 0.20 (means over ten trees). **c** L2 count relative to the truth over both compositions and depths (boxes, quartiles; lines, 10th–90th percentiles; shaded, ±25%); numbers, fraction of estimates within 25% of the truth. **d** Estimated relative to true L2 stem-cell mutation rate (posterior-weighted counts; pooled over tips and trees; 95% intervals from resampling trees), single leaves (per-organ model) and three leaves (replicate-group model); grey band, ±10%.

Using the same simulated trees, we then tested whether these counts recovered the known stem-cell mutation rate under neutrality and a stem-cell population at steady state. Posterior-weighted L2 counts from three leaves gave estimates of 0.95–1.00 times the true rate in most settings. At an L1 share of 0.20 and 62×, the estimate was lower, at 0.77. Single-leaf estimates were generally 20–36% above the true rate, except in that same setting, where the ratio was 1.01 (Fig. 6d; File S2, Table S8). The excess in single-leaf estimates reflected both organ sampling and errors in estimating fixed-mutation counts (File S1, Section S6).

We next asked whether the recovery seen in simulations extended to real sequencing data. Apricot fruit samples provided layer-enriched reference calls for testing recovery after candidate calling, and leaf samples allowed us to examine whether calls from one organ were supported in another from the same branch.

### Application to apricot sequencing data

#### Apricot data and testing strategy

We reanalysed the fruit and leaf sequencing data of Goel et al. (2024), who sampled seven branches of one apricot tree (Fig. 7a). Fruit peel is predominantly L1-derived and flesh L2-derived. These layer-enriched samples provided reference calls and allowed us to vary tissue composition by mixing reads. The leaf data allowed us to assess detection in conventional bulk tissue and check whether leaf calls were supported in fruit from the same branch.

**Figure 7:**
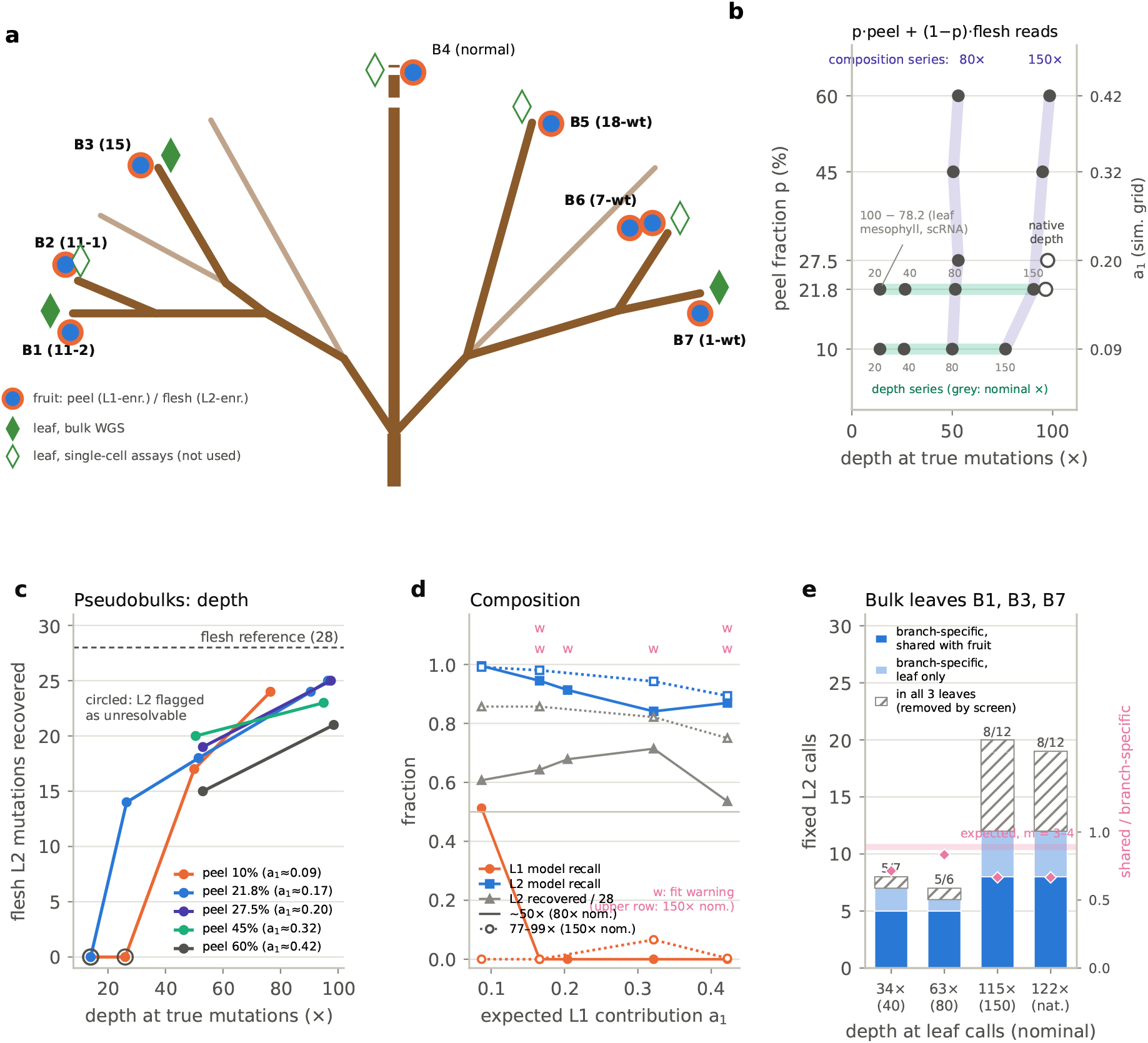
Apricot fruit pseudobulks and bulk leaves. (data of Goel et al. (2024); `vafSOMA` 0.4.2, per-organ model with the all-organs screen, B4 as normal). **a** Sampled branches, topology as in Johannes (2025b), redrawn after Goel et al. (2024) (CC BY 4.0), with the sample names of Goel et al. in parentheses; schematic, not to scale. Each branch contributed a fruit, separated into peel (L1-enriched) and flesh (L2-enriched), and a leaf; two adjacent fruits were used for B6. Bulk sequencing data of leaves were available for B1, B3 and B7. **b** Pseudobulk designs: peel fraction *p* (right axis, the corresponding L1 contribution *a*_1_ of the simulation grid) and depth at the reference mutations, with nominal depths in grey. Composition series at 80× and 150× nominal cross depth series at *p* = 10% and 21.8%; open circles, full available depth. **c** Reference L2 mutations (fixed in the flesh, *n* = 28) recovered in the pseudobulks; circles, designs in which L2 was flagged as unresolvable (no L2 calls). **d** Composition series at about 50× (nominal 80×; filled symbols) and 77–99 (nominal 150×; open symbols): model-implied recall of L1 and L2 and fraction of reference L2 mutations recovered; w, fit warning (upper row, nominal 150). **e** Fixed L2 calls in the leaves by depth at the calls (nominal depth in parentheses): branch-specific calls shared or not shared with the fruit of the same branch, and, hatched, the calls with alternative reads in all three leaves that the analysis without the screen adds (potential artefacts); numbers and diamonds (right axis), shared fraction of branch-specific calls; band, expected fraction for three to four stem cells (simSOMA).

#### Variant detection in fruit pseudobulks

We constructed fruit pseudobulks by downsampling and merging peel and flesh reads from the same branch. Their layer contributions were estimated from the

VAFs of layer-specific mutations: peel was about 69% L1-derived and flesh about 93% L2-derived (Materials and Methods). Changing the ratio of peel to flesh reads altered composition, while downsampling changed sequencing depth.

Analysing the flesh samples with `vafSOMA` as L2-enriched data provided 28 fixed L2 reference calls across the six branches other than the matched normal. We assessed their recovery across mixtures spanning the simulated L1 compositions and nominal depths from 20 to about 200× (Fig. 7b). This design followed the same underlying mutations through changes in composition and depth while retaining the sequencing and mapping properties of the original reads. Each mixture passed through joint Mutect2 candidate calling, `vafSOMA` fitting and posterior-weighted counting. After Mutect2 excluded low-mapping-quality reads, depth at the reference sites was 50–70% of nominal depth. All depths below refer to this post-filtering depth.

At about 97× and L1 shares up to 0.20, `vafSOMA` recovered 25 of the 28 L2 reference mutations and made three to six additional calls. At about 50×, recovery ranged from 15 to 20 reference mutations across compositions (Fig. 7c,d). At the lowest depths, 14–27×, at most 14 were recovered, and L2 was flagged as unresolvable in three of the four designs (File S2, Table S9).

L1 remained unresolvable in every pseudobulk of at least 50×, including L1 shares of 0.32 and 0.42, where simulations predicted partial resolvability. Increasing depth to about 95× at these two compositions did not resolve L1. Instead, some mutations fixed in L1 were assigned to L2: five of 11 additional L2 calls at an L1 share of 0.32, and 11 of 20 at 0.42, were L1 reference mutations from the peel. The corresponding numbers of recovered L2 reference mutations were 23 and 21. Of these two designs, only the latter raised a fit warning (File S2, Table S9). Two features of the apricot data can explain why L1 was harder to resolve than in the simulations. First, few L1 mutations were available to define an L1 component: the peel yielded 19 fixed L1 reference calls across the six branches, fewer than the 28 in L2, whereas in the simulations L1 carried about two thirds of the fixed L1 and L2 mutations. Second, the peels differed in composition among branches, and the peel VAFs of the L1 mutations assigned to L2 were higher than expected for heterozygous mutations at an L1 share of 0.69 (File S1, Section S8). In the pseudobulks of these branches, the L1 mutations therefore reached VAFs in the range of the L2 component. In simulations, variation in composition among organs likewise inflated L2 counts where L1 and L2 were both resolvable, without triggering a fit warning (Fig. 5c).

The two full-depth designs yielded posterior-weighted L2 rates of 1.22 × 10^*−*9^ and 1.45 × 10^*−*9^ per site and haploid genome per year. Our simulations suggest that single-organ estimates can exceed the targeted stem-cell mutation rate. Suitable replicate organs formed close in time were unavailable to test this prediction directly in apricot.

#### Variant detection in bulk leaves

We next analysed conventional bulk-leaf sequencing data from three branches. The assumed layer contributions, L1 0.13 and L2 0.84, were estimated from published leaf measurements and apricot single-cell RNA data (Materials and Methods). L1 was again unresolvable and L2 was resolvable. At the highest depths, `vafSOMA` called 12 fixed L2 mutations, all branch-specific (Fig. 7e; File S2, Table S10).

The sampling topology allowed us to apply the all-organs screen. Without it, analyses produced seven or eight additional L2 calls with alternative reads in all three leaves. At four investigated sites, alternative-allele reads were also present in fruit samples, including the matched normal, when low-mapping-quality reads were included, consistent with residual mapping artefacts (File S1, Section S8). Without the screen, the estimated leaf L2 rate per metre was 2.1–2.5 times higher (File S2, Table S10).

Eight of the 12 branch-specific calls were present in fruit from the same branch. Leaf and fruit can inherit different stem-cell lineages from their shared branch meristem, so concordance provides a check on fixation inferred from one organ. Simulations with three or four stem cells predicted 87–91% sharing. The observed sharing was lower (lower-tail binomial *P* = 0.06 for three and *P* = 0.02 for four stem cells). Missed fruit calls are unlikely to explain this. We found that, at three of the four leaf calls absent from the fruit, none of up to 166 reads per fruit design carried the alternative allele, and a fruit formed up to a year before the leaf changed the prediction little (85–92%; File S1, Section S8). The difference may instead arise because some of these calls are mutations fixed in the leaf alone, for example because leaf founder pools are smaller than assumed in the simulations, or are not fully fixed in the leaf. With 12 calls, this evidence is limited, but it agrees with the simulations in showing that a call in a single organ does not establish fixation in the branch meristem.

The VAFs of these 12 calls also informed the assumed tissue composition. A heterozygous mutation fixed in L2 would have an expected VAF of 0.42 if L2 contributed 0.84 of the sequenced DNA. The observed VAFs were closer to 0.5 (medians 0.47–0.49), consistent with a larger L2 contribution to sequenced DNA than indicated by the cell-based estimates. The published leaf measurements and the single-cell data of Goel et al. (2024) describe the leaf in terms of cells (their numbers, volumes or cell-type proportions), whereas VAFs depend on each layer’s share of the extracted DNA. The two can differ if cell types differ in DNA content per cell, for example through endopolyploidy, or in DNA yield during extraction, and cell-type proportions in single-cell data also depend on how efficiently each cell type is released and captured. Cell-based compositions are therefore approximations of the DNA shares that `vafSOMA` requires, and the VAFs of confident calls provide a check on them. In apricot leaves, the calls were robust to this difference. Indeed, with an assumed L2 share of 0.94, `vafSOMA` made 10 calls at the two highest depths, all among the 12 of the main analysis (File S1, Section S8).

These analyses supported recovery of most dominant-layer reference mutations at sufficient depth, while minor-layer recovery remained unsupported in the empirical samples. This distinction should guide the choice of tissues and sequencing depth in studies of plant somatic mutations.

## Discussion

Our analyses show that interpreting bulk somatic sequencing requires knowing the DNA contribution of each tissue layer. Mutations fixed in a dominant layer can produce a distinct VAF signal, whereas those fixed in minor layers can overlap with non-fixed variants and artefacts. In our simulations, L1 contributions of 0.17 or less were unrecoverable at every tested depth, while contributions of 0.20–0.25 required the highest depth. In apricot, L1 remained unresolvable across all fruit mixtures and in leaves. Increasing depth improves read precision, but does not by itself distinguish fixed mutations from background variants at similar VAFs.

`vafSOMA` uses tissue composition to make the interpretation of these signals explicit. It assigns candidates to layers, provides posterior-weighted counts for mutation-rate estimation, assesses layer support and model fit, and jointly analyses replicate organs. Its calling accuracy was close to that of the best VAF cutoff chosen in hindsight, whose optimum varied from 0.08 to 0.40 with composition and depth. Part of its advantage over cutoffs as commonly applied came from the all-organs screen. When the same screen was applied to all methods, calling accuracy was similar. `vafSOMA` additionally identifies which layers support separate counts and provides quantitative diagnostics unavailable from a cutoff alone.

Organ sampling also affects confidence in fixation. A variant confined to part of an organ can have a VAF close to the expectation for layer fixation if it arose in one of the organ’s founder cells, especially if that founder pool is small, so a single-organ call does not establish fixation in the branch meristem. In our simulations, single-organ counts overstated meristem fixation by about 40%. Analysing two or three organs formed close in time reduced this excess and yielded rates close to the known stem-cell mutation rate in most settings. These rate comparisons assumed neutrality and a steady-state stem-cell population without stem-cell displacement. In apricot, a third of the branch-specific leaf calls lacked support in the corresponding fruit. This comparison checks calls across organs, but suitable closely spaced replicates were unavailable to validate replicate-based rate estimation directly.

These findings affect comparisons among published somatic mutation estimates. Where L2 dominates leaf DNA, bulk sequencing primarily captures mutations fixed in L2, while those fixed in L1 and L3 can remain undetected. If two or three layers accumulate mutations independently at comparable rates over similar times, counting only one captures roughly half or a third of their combined fixed-mutation input. This concerns the lineages represented by an estimate, even when the rate for the recovered layer is accurate. The missing contribution could be larger where L1 accumulates mutations faster than deeper layers, as indicated by layer-enriched studies (Goel et al., 2024; Amundson et al., 2025). Within the recovered layer, single-organ sampling, permissive cutoffs and residual artefacts can raise estimates.

The relevant layer also depends on the biological question. On the positive side, mutations assigned to L2 are of particular interest, because L2 contributes to gametes in many species: L2-specific counts provide a direct view of the somatic mutations that can be transmitted to the next generation. However, where a search for causal mutations targets L1, as for many fruit-skin colour bud sports in fruit trees (Foster and Aranzana, 2018), our results indicate that bulk sequencing of leaves or whole fruits is likely to miss such mutations.

These considerations suggest practical sampling and reporting recommendations. Where possible, tissues should be enriched for the layer of interest, and two or three organs formed close in time should be sampled per branch to strengthen support for shared fixation. Independent composition measurements, supported by the VAFs of confident calls, can help assess assumed layer contributions. Sequencing depth should be evaluated after the caller’s read filters. Candidates present in every sampled organ warrant investigation, with the all-organs screen applied where the sampling topology permits. Rates should specify the layer, fixation criterion, composition, phasing and ploidy.

Several limitations remain. Only diploids analysed against a haploid reference were evaluated, although `vafSOMA` also supports phased calls. Composition must be supplied externally, and the confidence intervals do not include uncertainty in composition or model fitting. Because the layer composition of an organ needs to be measured only once per species and organ type, libraries of organ compositions could serve as a basis for `vafSOMA` analyses; otherwise, approximate values from the literature can be used. For example, published leaf compositions of *Arabidopsis thaliana* were close to the composition estimated from apricot single-cell data (Pyke et al., 1991; Tolleter et al., 2024; Goel et al., 2024), although such cell-based values can differ from the layers’ shares of the sequenced DNA (see Results). Effective genome sizes do not correct for the caller’s sensitivity at each layer’s expected VAF, and near-fixed variants can remain indistinguishable from fixed mutations. Layer support and fit diagnostics therefore guide interpretation without guaranteeing unbiased counts. The apricot data also supplied the simulation calibration and the pseudobulk reference calls, so the empirical analysis tests recovery after candidate calling without providing an independent truth set. Broader evaluation across species, tissues and callers remains necessary.

Despite these limitations, `vafSOMA` provides capabilities for somatic variant calling in plants that current approaches lack: assignment of fixed mutations to layers, posterior-weighted counts for rate estimation, diagnostics of layer support and joint analysis of replicate organs. Compared with the fixed VAF cutoffs routinely used in the literature, it achieved higher calling accuracy than the cutoffs as commonly applied, similar accuracy when both used the all-organs screen, and more interpretable results, because it reports which layers the data can resolve.

## Materials and Methods

### Input data: candidate calling with Mutect2

`vafSOMA` takes as input, for each plant, one table of candidate sites per minimum-depth cutoff, with the reference and alternative read counts of every organ at every site, and the effective genome size for each cutoff. We produced these inputs from aligned, duplicate-marked reads (one BAM file per organ, including the matched normal). For each plant, all sequenced organs were analysed jointly in Mutect2’s multi-sample mode (GATK 4.6.2.0) (Benjamin et al., 2019; McKenna et al., 2010), so that allelic depths are reported for every organ at every candidate site, including organs without alternative reads. One or more organs of the same plant served as the matched normal (apricot: the fruit of branch B4). Because the normal is itself an early-diverging part of the plant, mutations that arose before its divergence are removed as germline, and the candidate set is restricted to mutations that postdate this divergence. Emission thresholds were set to zero, candidates were filtered with FilterMutectCalls with no minimum allele fraction, and biallelic single-nucleotide variants with `FILTER=PASS` were retained. For each depth cutoff *d*_min_, a site was retained only if every organ had a depth of at least *d*_min_, giving one input table per plant and cutoff. The effective genome size for each cutoff was the number of haploid reference positions at which every organ had a SAMtools depth of at least *d*_min_, counting reads with base quality ≥10 and mapping quality ≥20, the mapping-quality filter of Mutect2. Settings, read processing and the choice of Mutect2 over Strelka2 are given in File S1 (Section S8).

### Statistical model

#### Data

For tree *t* and minimum-depth cutoff *d*_min_, let organs be partitioned into user-defined replicate groups with members *i* = 1, …, *r*_*g*_; ungrouped organs and control organs form singleton groups, and *n*_*g*_ denotes the number of non-control groups. For candidate *j*, 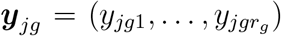 and ***d***_*jg*_ are the alternative read counts and depths of the members, zeros included. The model is fitted to the set of units

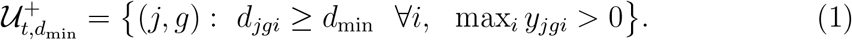

#### Components

Each unit has a latent state *Z*_*jg*_ ∈ *A*, where sh denotes shared background (one background VAF for all members of the unit), pr private background (an independent background VAF for each member) and 1, 2 and 3 fixation in L1, L2 or L3, with weights *π*_*a*_, ∑_*a*_ *π*_*a*_ = 1. The set of states is *A* = {sh, pr, 1, 2, 3} without sequencing error, extended by the error-only state err when *e >* 0 (below). With 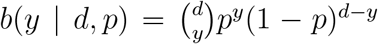 and the truncated exponential density

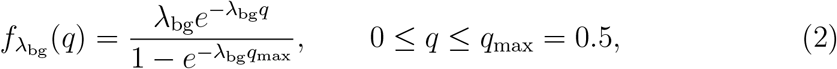

the unconditional component probabilities of a unit are

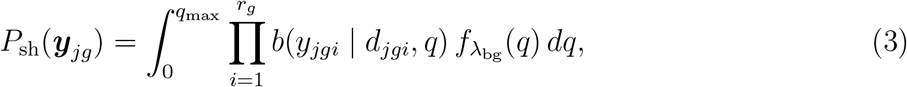

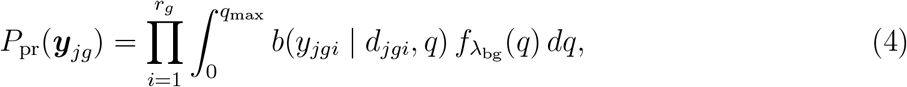

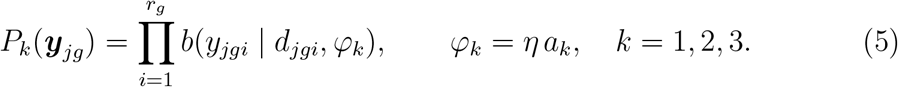

The shared component describes background that recurs in all replicates of a group, such as mapping artefacts; the private component describes variants confined to single organs. Integrals over *q* are evaluated by 96-point Gauss–Legendre quadrature. The layer components can optionally be beta-binomial with a shared overdispersion parameter *ψ*, the intra-class correlation of reads (File S1, Section S2).

#### Observation model of the reads

`vafSOMA` can use a read-level observation model whose parameters are calibrated for each data set (apricot: File S1, Section S7). A site with allele fraction *v* reports the alternative allele with probability *p*_*e*_(*v*) = *e* + (1 −2*e*)*v*, where *e* is a symmetric per-read sequencing error, and the allele fraction is overdispersed with intra-class correlation *ψ*. In Equations 3–5, *b*(*y* | *d, q*) is then evaluated at *p*_*e*_(*q*), and the layer components are beta-binomial with mean *p*_*e*_(*φ*_*k*_) and the matching dispersion (moment-matched; File S1, Section S2). With *e >* 0 the background also contains an error-only state, in which no replicate of the group carries the variant and every alternative read is an error,

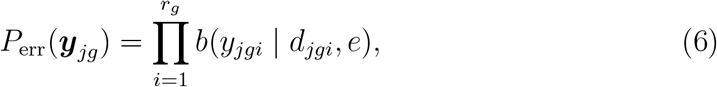

with its own weight *π*_err_, conditioned on a positive replicate like the other components. At a mutated site, organs of other groups show such stray error reads next to zeros in their sibling replicates; without this state only the exponential background with a very large *λ*_bg_ can represent them, and in simulations with *e* = 10^*−*3^ the replicate-group model then absorbed background into the layer components (File S1, Section S2). *e* and *ψ* are taken from the calibration (*e* = 3 × 10^*−*4^ and *ψ* = 1*/*141 for apricot) rather than estimated; with *e* = *ψ* = 0 the model is that of earlier versions. *π*_err_ counts as background in all outputs.

#### Conditional likelihood

Each component is conditioned on at least one positive replicate,

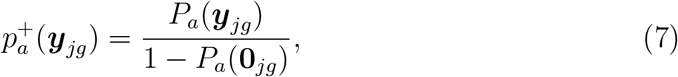

where **0**_*jg*_ is the all-zero vector at depths ***d***_*jg*_, and the log-likelihood is

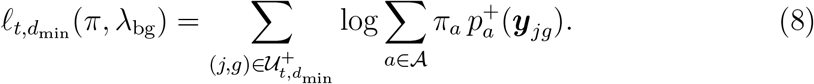

Conditioning on detection keeps the many sites without alternative reads from pulling the background toward zero. Including the zero counts of sibling replicates in the fixed components makes absence in a replicate informative. A mutation fixed in the major layer is unlikely to yield no reads in a replicate at moderate depth.

#### Per-organ case

If *r*_*g*_ = 1 for all groups, *P*_sh_ = *P*_pr_, and Equation 8 reduces exactly to the mixture of positive per-organ observations,

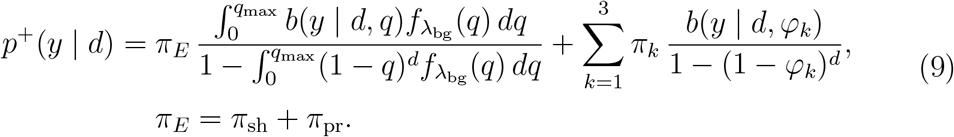

Equation 9 is written for *e* = *ψ* = 0. With the observation model, *b* is evaluated at *p*_*e*_(*q*) in the background term and the layer terms are beta-binomial with mean *p*_*e*_(*φ*_*k*_) (as above), and the error-only term *π*_err_ *b*(*y* | *d, e*)*/*{1 − (1 − *e*)^*d*^} is added; the background weight reported in the outputs is then *π*_*E*_ + *π*_err_. In the implementation, the per-organ code path is used when no groups are declared (File S1, Section S1).

#### Expected VAFs and target size

The layer proportions *a*_*k*_ are supplied by the user, and *η* is 1*/*2 for unphased and 1 for phased counting in a diploid. This read-level factor must not be confused with the factor 2 in the rate denominator (Equation 11), which is the diploid mutational target size and applies regardless of phasing (File S1, Section S1). For a layer-enriched sample, such as fruit flesh for L2, the user specifies the enriched layer and its purity *a*_*k*_ instead of the bulk proportions; only that layer’s component is fitted, with *φ*_*k*_ = *ηa*_*k*_. Where another layer contributes substantially, as in apricot peel (about 69% L1), the sample can instead be analysed as a bulk organ with its estimated composition.

### All-organs screen

Optionally, candidates with at least one alternative read in every sampled non-control organ, at the lowest depth cutoff, are removed from all cutoffs before the fit. The screen is valid only if no lineage after the divergence from the matched normal is shared by all sampled organs; it then removes recurrent artefacts (mapping, paralogs, library-wide errors), but also the rare true mutations that drifted to fixation in all sampled lineages while being lost from the lineage of the normal. `vafSOMA` requires a declared control organ and at least three non-control organs per tree. Removing these candidates before the fit, rather than from the calls afterwards, keeps them out of the fitted background: otherwise they make the background heavier, and true layer-fixed mutations are absorbed as background. All simulations and apricot analyses of this study used the screen, with B4 as control; the apricot analyses were repeated without it (File S1, Section S5). The screen is applied at the lowest depth cutoff of the input, which was the first cutoff of the depth window in the per-metre apricot analyses and the lowest available cutoff in the per-year analyses; within the window, the posterior-weighted L2 counts of the two were identical.

### Estimation

For fixed *λ*_bg_, the weights are estimated by expectation-maximisation. With responsibilities 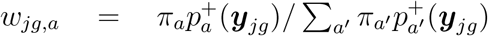, the update is 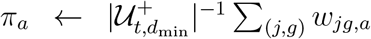, iterated to a relative change in *l* below 10^*−*8^ from multiple starting points (deterministic and random Dirichlet starts; 16 by default). Identical observation vectors are collapsed into weighted rows. The profile likelihood 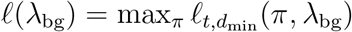 can be multimodal; it is maximised by a coarse log-spaced scan, exhaustive zoom refinement around the best points, a bounded local polish and a final multistart re-fit (File S1, Section S3). When layer proportions or overdispersion are given as small grids, fits are combined by likelihood weights, and a phased single-layer design (*φ*_*k*_ = 1) triggers an automatic grid just below 1 (File S1, Section S3). An optional sensitivity sweep refits the model under perturbed layer proportions and reports each result separately (File S1, Section S4).

### Posterior classification

The posterior probability of state *a* for unit (*j, g*) is

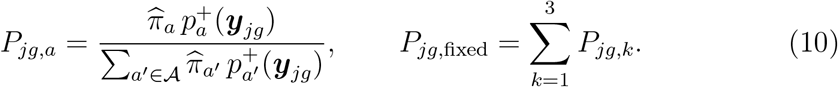

For non-redundant candidate lists, the likelihoods of a candidate’s groups are multiplied under a single state, 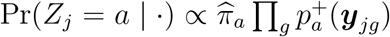, with the product over groups in which the candidate was observed. Over the selected depth window, a candidate is called fixed in layer *k* if its maximum-a-posteriori state is layer *k* at all cutoffs (a user-adjustable fraction, 1 by default), fixed with ambiguous layer if it is fixed at all cutoffs but the layer varies, background if it is never fixed, and unstable otherwise (File S1, Section S5).

### Fixed-mutation counts and mutation rates

For group *g* (an organ, or a replicate group) and cutoff *d*_min_, *N*_*g*_ = ∑_*j*_ *P*_*jg*,fixed_ is the posterior-weighted number of fixed mutations, and *N*_*g,k*_ = ∑_*j*_ *P*_*jg,k*_ that of layer *k*; groups without positive observations contribute zeros. `vafSOMA` reports these counts per group and cutoff, the single-observation posteriors behind them, the effective genome size 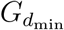 of each cutoff and the length *L*_*g*_ of each group (the mean of its members’ lengths), as well as hard-call (maximum-a-posteriori) counts. Rates are not computed by `vafSOMA`. We computed them from these outputs as the unweighted mean of per-group ratios,

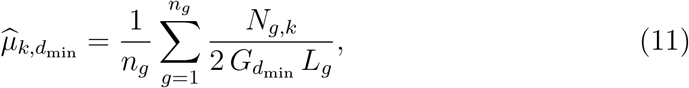

where *L*_*g*_ is measured from the divergence with the normal and *n*_*g*_ is the number of non-control groups. Averaging ratios rather than pooling counts treats each group as one replicate estimate, and 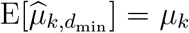 whenever 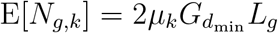 for each group, irrespective of covariance between groups due to shared ancestry (File S1, Section S6). All counts and rates reported in this study are these posterior-weighted quantities; counts and rates of layers flagged as unresolvable (see below) are not interpreted, because the posterior mass of such a layer comes largely from background candidates. Hard-call counts and rates differed from the posterior-weighted values by a few percent for the resolvable layers analysed here. Rates are reported per site and haploid genome and per unit of the supplied length *L*_*g*_ (per year if *L*_*g*_ is an age, per metre if it is a distance; both are reported for the apricot data). The rate of a window of cutoffs is the mean of 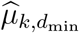 over its cutoffs. Because the depth tiers are nested subsets of the same sites and reads, confidence intervals are percentile intervals from a bootstrap (*B* = 1,000) that resamples the candidate variants of each group once and applies the same draw to every cutoff of the window, recomputing the window-averaged rate; the fitted weights and background are held fixed, so the intervals do not include fitting uncertainty (File S1, Section S6). The calculation is provided as a script (Software and data availability). What this rate estimates, and under which assumptions, is derived in Appendix A.

### Depth-window selection

The analysis is repeated over a range of minimum-depth cutoffs, and `vafSOMA` summarises calls and counts over a contiguous window of cutoffs in which they are stable. The window is chosen from the density of hard-called fixed mutations, 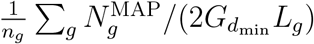, which removes the change of the callable genome with the cutoff: the window starts at a cutoff of at least 12, comprises at least four cutoffs, and has a coefficient of variation ≤ 0.15 and a maximum relative change between adjacent cutoffs ≤ 0.25; among the windows that pass in all trees of an analysis, the one with the most cutoffs is chosen (ties broken by the smaller coefficient of variation), and if none passes, all cutoffs are used. These criteria detect jumps but not a gradual trend, so counts and rates should also be inspected as a function of *d*_min_ (File S3, Figure S3).

### Post-fit diagnostics

Two diagnostics are computed from the fitted parameters and the input data; they do not change fits, posteriors or calls. *Layer resolvability*. For each tree, depth cutoff and active layer *k*, the model-implied recall is the probability that an observation of a mutation fixed in layer *k*, with alternative reads *y* ~ *b*(· |*d, φ*_*k*_) conditional on *y* ≥ 1, has maximum-a-posteriori state *k* under the fitted mixture, averaged over the empirical depth distribution of the observations. Averaged over the selected depth window, a recall below 0.5 marks the layer as unresolvable; its fixed mutations would mostly be absorbed by the background or by another layer, and its separate layer-specific count and rate are not reported. A layer is also marked as unresolvable if its expected VAF lies within the fitted background, 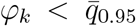, where 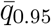 is the 95% quantile of the fitted truncated exponential (Equation 2), 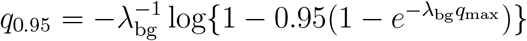, averaged over the cutoffs of the window. This condition was added because, with a light background and a small minor-layer share, the minor-layer component can absorb background candidates and reach a high model-implied recall without a fit warning (Appendix B). In simulations with the calibrated observation model, the model-implied recall of resolvable layers exceeded the recall of truly fixed mutations by 0.02 on average, but by −0.09 to +0.08 between settings (File S1, Section S5); it is therefore used only to flag unresolvable layers, and counts are not corrected by it. *VAF fit check*. For each depth cutoff, positive observations are binned by VAF (bin edges 0, 0.025, 0.05, 0.075, 0.10, 0.15, 0.20, 0.25, 0.30, 0.40, 0.50, 1), and the observed number per bin is compared with the expected number under the fitted mixture, ∑_*i*_ Pr(*y*_*i*_*/d*_*i*_ ∈ bin | *d*_*i*_, *y*_*i*_ ≥ 1), by Pearson residuals. A bin with an expected count of at least 10 and a residual of at least 4 in absolute value is flagged, and a fit warning is raised when flagged bins occur in at least half of the depth cutoffs of the selected window.

### Simulations

#### Trees

Somatic evolution was simulated with simSOMA 0.2.3 (Johannes, 2026a,b) on an approximate topology of the apricot tree studied by Goel et al. (2024), scaled to a mean root-to-tip length of 11 years, with seven sampled branch tips. Each meristem layer was simulated as an independent cell lineage, with mutation rates of 0.95 (L1) and 0.55 (L2, L3) new mutations per diploid stem-cell genome and year and shared developmental parameters (*m* = 3 apical stem cells, no stem-cell displacement). Unless stated otherwise, a leaf and a fruit were sampled at each tip, and each setting comprised ten trees. For each mutation and organ, the simulation yields the carrier-cell fraction *f*, the fraction of the organ’s cells in the mutation’s layer that carry it; the bulk VAF of a mutation of layer *k* is 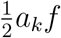 (unphased diploid). Parameters are listed in File S1 (Section S8).

#### Observation model

Reads were generated with the observation model calibrated on the corresponding apricot design (File S1, Section S7): the per-sample mean depths of the real design, log-normal site and site-by-sample depth factors, beta-binomial reads (concentration 140) with a symmetric sequencing error of 3 × 10^*−*4^, background candidates resampled from the real non-somatic candidates of the same design, the calling edge of the real design and Mutect2’s normal filter. From these background pools we removed 193 sites with evidence of real somatic mutations in the layer-enriched apricot samples (called fixed in the flesh or peel, or with at least three alternative reads and a VAF of at least 0.15 there and no alternative read in B4). Fruit designs comprised the six non-normal fruits at full depth and at 150, 80 and 40× nominal (103, 84, 46 and 26× at true mutations), and leaf designs the leaves of B1, B3 and B7, each with the fruit of B4 as matched normal. `vafSOMA` 0.4.2 was run with the calibrated read model (*e* = 3 × 10^*−*4^, *ψ* = 1*/*141), with L1 and L2 modelled (L3 not modelled, as in the apricot analysis: with the empirical background, an L3 component at *φ*_3_ ≈ 0.04 absorbed the background), with the all-organs screen, and with the window fixed to four nested depth tiers.

#### Settings

Layer composition: *a*_3_ = 0.075, *a*_2_ = 0.925 − *a*_1_. The cutoff comparison used *a*_1_ ∈ {0.087, 0.204, 0.322, 0.422} and the composition sweep eight further values from 0.05 to 0.45. The stress tests replaced the resampled background by a parametric one (2,000 sites, gamma with shape 2 and mean 0.023), doubled the background pool, added a heavy background component (exponential with mean 0.06, at 0.25 or 1.0 times the number of pool sites), lowered the read concentration to 50 or 20, raised the sequencing error to 10^*−*3^, or supplied an L1 contribution of 1.25*a*_1_ or 0.75*a*_1_ to `vafSOMA` while simulating with *a*_1_. Organ-specific composition: 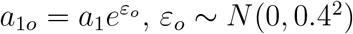, at full depth and *a*_1_ ∈ {0.204, 0.322}, fitted with the nominal composition, the true organ-specific compositions, or compositions estimated from the VAFs of first-pass L2 calls.

#### Replicate organs

Three leaves were formed at each branch tip, 0.02 years apart. The same trees were analysed with one, two or three leaves per tip (*a*_1_ ∈ {0.087, 0.204}; leaf designs at native depth and 80× nominal, 108 and 62× at true mutations) with the per-organ model, the per-organ model followed by a concordance screen (a fixed call is kept if made in at least two leaves of the tip), and the replicate-group model with the leaves of each tip as a group. Background candidates were drawn in profile mode, which applies each real candidate’s allele fraction to all leaves and draws the reads of each leaf; the all-organs screen removed 29–49% of the candidates in these designs, so that background remained in the fit. For the rate check, posterior-weighted L2 counts were compared with the true rate per tip, 0.55 *q* (*L* −*L*_0_), where *q* is the fraction of the tree’s fixed L2 mutations that entered the input tables and *L* − *L*_0_ the age since divergence from B4 (File S1, Section S8).

#### Truth and scoring

A mutation was scored as fixed in an organ if its carrier fraction satisfied 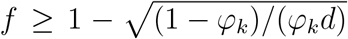 at the observed depth *d*, that is, if its expected VAF was within one binomial standard deviation of that of a fixed mutation; strict fixation (*f* ≥ 0.999) is reported in File S1. For the replicate analysis, fixation in the branch meristem required fixation in all three leaves of the tip. Calls were scored per site and organ, and layer-specific sensitivity, precision and F1 score used the layer assigned by `vafSOMA`. For the comparison with fixed cutoffs, all calls were scored jointly, mutations fixed in L3 were excluded, and the F1 score was compared with the maximum over cutoffs from 0.02 to 0.40 in steps of 0.01 in the same data set. A cutoff called an organ-level candidate with VAF at least *t* and at least three alternative reads; the composition-aware cutoff used *t* = (0.02 + *φ*_2_)*/*2. Cutoffs were evaluated as commonly applied and after the all-organs screen. `vafSOMA` counts are posterior-weighted (sums of *P*_*jg,k*_) for layers not flagged as unresolvable.

### Apricot data

#### Fruit pseudobulks

Reads from the peel and flesh of fruits from seven branches of one apricot tree (*Prunus armeniaca* cv. Rojo Pasión) (Goel et al., 2024) were aligned to the Currot haplotype. For each branch, peel and flesh BAM files were downsampled to the target peel fraction *p* and total depth *D* and merged. The designs were *p* ∈ {10, 21.8, 27.5, 45, 60%} at 80×, *p* ∈ {10, 21.8%} at 20 and 40×, *p* ∈ {10, 21.8, 45, 60%} at 150×, and *p* ∈ {21.8, 27.5%} at the full depth (116– 227×). Each design was called with Mutect2 as above, with the B4 pseudobulk as normal. In Goel et al.’s data, the peel is about 69% L1-derived and the flesh about 93% L2-derived, so that the L1 share of a pseudobulk is approximately 0.02+0.67*p* and its L2 share 0.93 − 0.62*p*; the remainder is L3, which was not modelled. Reference calls were obtained by running `vafSOMA` on the flesh (L2 purity 0.93) and peel samples as layer-enriched data; a reference mutation was counted as recovered if it was called fixed in L2 in the pseudobulk of the same branch. Pseudobulks were analysed with `vafSOMA` 0.4.2 (per-organ model, all-organs screen, B4 as control, eight EM starts). As a depth measure, we used the median Mutect2 depth of the non-normal organs at the reference L2 mutations present in each design (13– 26 mutations). Rates per metre used the measured distances from the divergence with B4 to the sampled tip (3.18–3.68 m), and rates per year the ages of the branch segments of Johannes (2025b, Fig. 5a) from the crown base, where B4 diverges, to each tip (10.0–11.6 years), with effective genome sizes at mapping quality ≥ 20. Details, including the estimation of tissue compositions and the read-level checks, are given in File S1 (Section S8).

#### Bulk leaves

Leaves of branches B1, B3 and B7 were analysed at their native depth (154–163×) and downsampled to 40, 80 and 150×, jointly called with Mutect2 against the B4 pseudobulk of the full-depth design with *p* = 21.8%, and analysed with `vafSOMA` 0.4.2 as above. The leaf composition was set to the mean of three published estimates, normalised to the three layers (L1 0.13, L2 0.84; L3 not modelled) (Pyke et al., 1991; Tolleter et al., 2024; Goel et al., 2024), and analyses were repeated with L1 0.03 and L2 0.94. Branch-specific fixed L2 calls were classed as shared if the variant was a fixed L2 call in the flesh of the same branch or had at least three alternative reads and a VAF of at least 0.15 in that branch’s flesh or fruit pseudobulks. The expected sharing was obtained from simSOMA simulations of leaf and fruit at the same branch tip (File S1, Section S8).

### Software

`vafSOMA` (version 0.4.3) is implemented in Python (≥3.10) with NumPy, pandas and SciPy, and is available at https://github.com/jlab-code/vafSOMA under the MIT licence. The analyses used version 0.4.2, which added the all-organs screen and the background condition of the resolvability flag to the observation model of version 0.4.0; without these options, its fits, posteriors and calls are those of version 0.4.0. Version 0.4.3 changes only the documentation and gives identical results. Mutation rates and their bootstrap intervals were computed from the outputs of `vafSOMA` with a script (`apricot_rates_v042.py`, in the data record below). The observation model used for the simulations and its apricot calibration are described in File S1 (Section S7); its code is available from the corresponding author upon request. Simulation and analysis scripts and the list of sites excluded from the background pools are provided on Zenodo (https://doi.org/10.5281/zenodo.23884962). Large language models were used as follows. ChatGPT (OpenAI; GPT-6.1 Sol) was used for language editing of the manuscript text, with prompts asking for clearer and more concise wording of paragraphs written by the authors. Claude (Anthropic; Claude Opus 5.5), accessed through claude.ai, was used to write and run analysis and figure code, to implement parts of `vafSOMA`, and to edit the text. The authors checked all code, results and text and take responsibility for them.

## Data availability

The `vafSOMA` source code is available at https://github.com/jlab-code/vafSOMA (MIT licence; version 0.4.3 archived at https://doi.org/10.5281/zenodo.23882759), with a tutorial at https://github.com/jlab-code/vafSOMA-tutorial (archived at https://doi.org/10.5281/zenodo.23884858). The apricot sequencing data of Goel et al. are available from ENA (PRJEB71142). The derived pseudobulk and leaf VAF tables, effective genome sizes, `vafSOMA` outputs and the per-data-set simulation results are provided on Zenodo (https://doi.org/10.5281/zenodo.23884962), together with the analysis and simulation scripts. Supplementary material is available online: File S1 (Supplementary Text) contains the model details, numerical estimation, posterior classification and diagnostics, rate calculation and bootstrap (Sections S1–S6), the simulation of layered bulk data (Section S7) and detailed methods (Section S8); File S2 contains Supplementary Tables S1–S10, and File S3 Supplementary Figures S1–S3.

## Appendix A: What the rate measures

A mutation is fixed in a sampled organ if it lies on the cell lineage ancestral to all cells of the organ’s layer, and fixed in the branch meristem if it lies on the lineage ancestral to all of the meristem’s stem cells; the latter is approached by requiring fixation in all replicate organs of a branch tip. Mutations that arose before the sampled lineage diverged from the matched normal are removed as germline. For neutral mutations and a stem-cell population at steady state, the expected numbers of fixed mutations of layer *k* in a sample at length *L* are

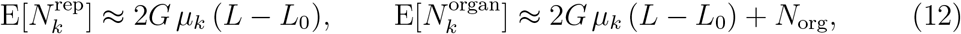

where *µ*_*k*_ is the mutation rate of the apical stem cells per site and haploid genome per unit time, *L*_0_ the part of the path before the divergence from the normal, and *N*_org_ ≥ 0 the number of mutations fixed in the organ because they were carried by its few founder cells without being fixed among the stem cells. For replicate-level fixation, three contributions combine: mutations that arose after the divergence from the normal and became fixed among the stem cells before the organs formed, which fall short of 2*Gµ*_*k*_(*L* − *L*_0_) because the most recent ones are not yet fixed; mutations that arose before the divergence, were still segregating among the stem cells when the branch split from the normal, and became fixed in the branch alone; and mutations that arose in the divisions forming a new branch meristem and became fixed through the founding bottleneck. Their relative contributions depend on the developmental parameters. In our simulations (no stem-cell displacement; Materials and Methods), replicate-level counts from two or three organs fell 3–7% below 2*Gµ*_*k*_(*L* − *L*_0_) on average, and the three-leaf ratio ranged from 0.72 to 1.26 among branch tips (File S1, Section S6). The balance therefore depends on the developmental parameters and the topology, and 2*Gµ*_*k*_(*L* − *L*_0_) is an approximation. Under the idealisation of Johannes (2025a), in which stem-cell mutations become fixed at the next bottleneck and transit-cell mutations are rare, *N*_org_ vanishes. 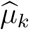 of a resolvable layer therefore estimates *µ*_*k*_ when computed from replicate-level calls and with *L*_*g*_ an age measured from the divergence with the normal; with *L*_*g*_ measured from the base of the plant, it is biased downward by the mean fraction of the path before the divergence, and rates per metre estimate *µ*_*k*_ only if growth is uniform in time. Mutations shared by a clade of sampled branches are counted once in each of its groups, which leaves the expectation unchanged.

## Appendix B: Background condition for layer resolvability

The model-implied recall of layer *k* is computed under the fitted mixture. If the layer component absorbs background candidates, the fitted weight 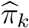 grows, and with it the posterior of state *k* at VAFs near *φ*_*k*_, so that the recall can be high although few candidates of the component are mutations fixed in layer *k*. This happens when *φ*_*k*_ lies inside the background, that is, when the background density at *φ*_*k*_ is not small compared with its mass at lower VAFs. For the truncated exponential background (Equation 2), the *p*-quantile of the background VAF is

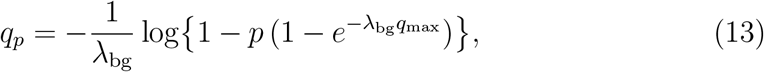

which tends to − log(1 − *p*)*/λ*_bg_ for *λ*_bg_*q*_max_ ≫ 1 and to *p q*_max_ for a flat background. A layer with *φ*_*k*_ *< q*_0.95_, averaged over the cutoffs of the window, is reported as unresolvable irrespective of its recall. In the simulations, this condition flagged every data set in which L1 had been reported resolvable with a count far above the truth (parametric background at *a*_1_ = 0.087; resampled background at *a*_1_ = 0.05; after removing shared candidates at *a*_1_ = 0.087; 36 data sets), and none in which L1 was correctly resolvable (57 data sets). In the apricot data it flagged L1 in the pseudobulk design with 10% peel at about 50×, where L1 had been reported resolvable without support from the peel references (*φ*_1_ = 0.044 against *q*_0.95_ = 0.054), and no L2 in any design. The value 0.95 is a default; with `resolvability_background_quantile: none` the flag reverts to the recall rule alone.

